# Non-cell autonomous control of presynaptic remodeling by the hypothalamic autophagy/NPY axis

**DOI:** 10.1101/2025.06.20.660653

**Authors:** Giovanna Cazzolla, David Toppe, Gina Krause, Gaga Kochlamazashvili, Janine Lützkendorf, Ina M. Schedina, Yannic Kerkhoff, Eric Reifenstein, Max von Kleist, Herbert Herzog, Anne Albrecht, Dietmar Schmitz, Volker Haucke, Stephan J. Sigrist, Marta Maglione

## Abstract

Macroautophagy/autophagy, a critical cellular degradation pathway essential for maintaining neuronal proteostasis, declines with age and has been increasingly implicated in the regulation of synaptic integrity and circuit resilience. Neuropeptide Y (NPY), the most abundantly expressed neuropeptide in the mammalian brain, has emerged as a key modulator of both autophagy and aging-related processes. In *Drosophila*, the NPY-family peptide short Neuropeptide F (sNPF) has been shown to causally influence aging-associated changes in synaptic architecture and function, particularly at the presynaptic active zone (AZ), via non-cell autonomous mechanisms.

Extending this concept to mammals, we investigated whether NPY and autophagy interact within NPY-secreting neurons to regulate age-related AZ remodeling. Our results indicate that hypothalamic NPY/AgRP neurons may exert geroprotective effects through the release of NPY and potentially other signaling molecules, thereby influencing both metabolic homeostasis and brain-wide synaptic function. These data suggest a conserved role for autophagy in maintaining presynaptic organization and resilience during aging.

## Introduction

Autophagy is a conserved lysosomal degradation pathway essential for maintaining neuronal proteostasis and synaptic function. Postmitotic neurons critically depend on autophagy to sustain long-term viability and circuit stability[1]. With age, autophagy declines, contributing to synaptic dysfunction, cognitive decline, and neurodegeneration[2–4]. However, how neuronal autophagy intersects with brain resilience and aging-related synaptic remodeling remains incompletely understood.

In *Drosophila*, we previously identified a presynaptic remodeling program termed PreScale, triggered by aging, oxidative stress, and sleep deprivation[5–8]. PreScale is marked by enlargement of the presynaptic active zone (AZ) and increased levels of core components such as Bruchpilot (BRP)[5]. Mimicking PreScale by increasing *brp* gene dosage promotes sleep homeostasis and longevity[7]. Importantly, spermidine (Spd), a natural autophagy inducer[9], prevents PreScale-associated memory decline, indicating a link between autophagy and presynaptic integrity[5,6].

Autophagy impairment in higher *Drosophila* brain centers, particularly the Mushroom Body (MB), can trigger PreScale non-cell autonomously via reduced neuropeptide short Neuropeptide F (sNPF), revealing a systemic neuropeptide-autophagy interaction[10]. In mammals, the sNPF homolog Neuropeptide Y (NPY), mainly produced in the hypothalamus, similarly promotes autophagy and longevity and declines with age[11–13]. NPY/AgRP neurons regulate energy homeostasis and are emerging as central integrators of aging and metabolic control[12,13].

Here, we test the hypothesis that autophagy within hypothalamic NPY/AgRP neurons regulates presynaptic remodeling in distant brain regions, bridging metabolic state with synaptic resilience. We further explore whether dietary spermidine supplementation (Spd-S) protects this axis in aged mice.

## Results

### Autophagy in hypothalamic NPY/AgRP secreting neurons maintains NPY non-cell autonomously

Previous studies demonstrated a bidirectional interaction between neuronal autophagy and NPY in mice. Here, NPY is required for autophagy induction by caloric restriction in hypothalamic and cortical neurons[12,14], while the 5’ adenosine monophosphate-activated protein kinase (AMPK) controlled autophagic status of hypothalamic NPY/AgRP secreting neurons modulates NPY expression at transcriptional level[15]. To further address whether autophagy in hypothalamic NPY/AgRP neurons would be required to maintain proper NPY signaling *in vivo*, we generated mice lacking the autophagy regulator ATG5 selectively in these neurons (*atg5*^AgRP^ cKO) by crossing *atg5*^flox/flox^ mice[16,17] to AgRP-IRES-Cre mice[18], a previously described mouse line expressing the Cre recombinase selectively in hypothalamic NPY/AgRP neurons[18–20]. *atg5*^AgRP^ cKO mice were born at mendelian ratio and their body weight did not differ from isogenic controls between postnatal day P40-P60 (Fig. S1A-C). At this developmental stage, AgRP-Cre-mediated deletion of *atg5* resulted in a marked reduction of ATG5 protein levels within NPY/AgRP neurons (Fig. S1D,E).

Concurrently, the autophagy cargo receptor SQSTM1/p62 (sequestosome 1) (Figure 1A,B) and the autophagosomal marker LC3 (Fig. S1F,G) were elevated in hypothalamic NPY/AgRP neurons, collectively indicating the expected impairment of autophagy. Notably, a modest but statistically significant decrease in NPY peptide levels was observed in the hypothalamic NPY/AgRP neurons of *atg5*^AgRP^ cKO (Figure 1C,D). Surprisingly, hippocampal hilar NPY-secreting neurons showed an even greater decrease in NPY signal in *atg5*^AgRP^ cKO (Figure 1E,F), accompanied by elevated SQSTM1/p62 levels (Figure 1G,H). In contrast, expression levels of ATG5, SQSTM1/p62 and LC3 in hippocampal granule cells remained unchanged following *atg5*^AgRP^ cKO (Fig. S1H-M), suggesting an impact specifically on NPY-secreting neurons beyond the hypothalamus.

**Figure 1.**
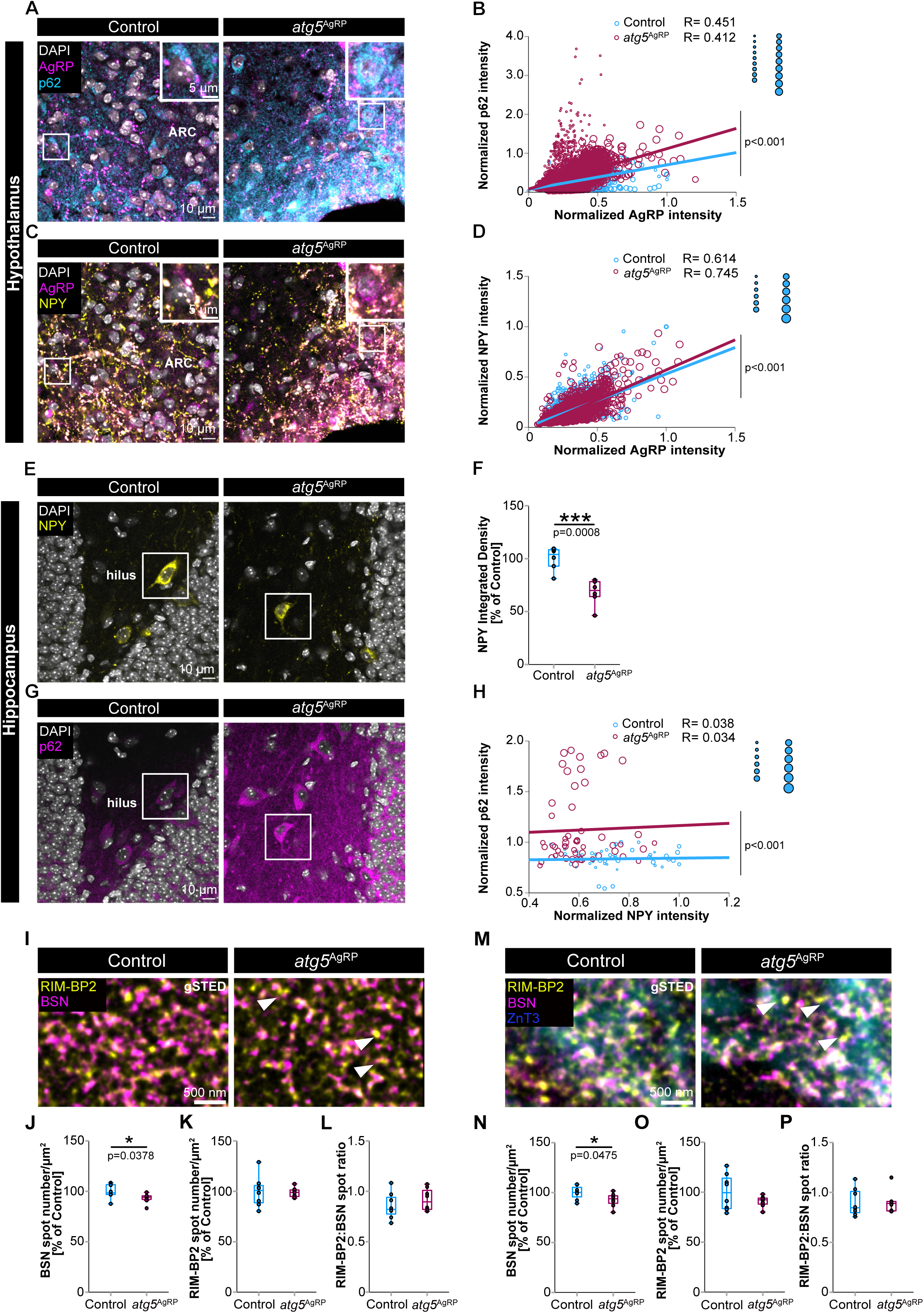
Autophagy in hypothalamic NPY/AgRP neurons controls NPY and presynaptic remodeling non cell autonomously. (**A**) Representative confocal images of AgRP and SQSTM1/p62 immunoreactivity in the hypothalamic ARC of control and *atg5*^AgRP^ cKO mice at P40-60. Scale bars: 10 µm, insets: 5 µm. **B**) SQSTM1/p62-AgRP mean intensity analysis in the ARC showed selective autophagy cargo receptor SQSTM1/p62 accumulation in hypothalamic NPY/AgRP neurons of *atg5*^AgRP^ cKO mice compared to controls. SQSTM1/p62 levels were related to AgRP levels with a GLMM with gamma regression (p < 0.001; control: *n =* 3208 cells; *atg5*^AgRP^ cKO: *n =* 3555 cells; *N =* 9 mice/genotype). **C**) Hypothalamic NPY/AgRP neurons immunostained for AgRP and NPY in the hypothalamic ARC of control and *atg5*^AgRP^ cKO mice at P40-60. Scale bars: 10 µm, insets: 5 µm. **D**) *atg5*^AgRP^ cKO resulted in significantly decreased NPY levels in NPY/AgRP neurons. NPY levels were related to AgRP levels with a GLMM with gamma regression (p < 0.001; control: *n =* 2655 cells; *atg5*^AgRP^ cKO: *n* = 2904; *N =* 6 mice/genotype). **E**) NPY immunolabelling in the dentate gyrus hilar region within the hippocampus of control and *atg5*^AgRP^ cKO mice. Scale bars: 10 µm. **F**) NPY integrated density showing impaired hilar NPY levels in *atg5*^AgRP^ cKO mice (68.08 ± 5.03 %, *n =* 6 mice) compared to controls (100 ± 4.53 %, *n =* 6 mice) (p = 0.0008; Student’s t test). **G**) SQSTM1/p62 immunolabelling in the dentate gyrus hilar region within the hippocampus of control and *atg5*^AgRP^ cKO mice at P40-60. Scale bars: 10 µm. **H**) SQSTM1/p62-NPY mean intensity analysis, *atg5*^AgRP^ cKO lead to SQSTM1/p62 accumulation in hippocampal NPY neurons. SQSTM1/p62 levels were related to NPY levels with a GLMM with gamma regression (p < 0.001; control: *n =* 47 cells; *atg5*^AgRP^ cKO: *n =* 54 cells; *N =* 6 mice/genotype). **I**) Time-gated STED (gSTED) microscopy of RIM-BP2 and BSN at hippocampal CA3-CA1 synapses in control and *atg5*^AgRP^ cKO mice at P40-60. Scale bars: 500 nm. **J-L**) Quantification of BSN (J; control: 100 ± 2.46 %; *atg5*^AgRP^ cKO: 93.2 ± 1.65 %; *n =* 8 mice), RIM-BP2 (K; control: 100 ± 5.30 %; *atg5*^AgRP^ cKO: 98.88 ± 1.62 %; *n =* 8 mice) spot density and RIM-BP2:BSN spot ratio (L; control: 0.85 ± 0.05; *atg5*^AgRP^ cKO: 0.92 ± 0.04; *n =*8 mice) showed a significant decrease in BSN spot number/µm^2^ in *atg5*^AgRP^ cKO mice compared to controls (BSN spot density: p = 0.0378; *n =* 8 mice/genotype; Student’s t test). **M**) Time-gated STED microscopy of RIM-BP2 and BSN at hippocampal MF-CA3 synapses identified by their marked ZnT3 expression (confocal) in control and *atg5*^AgRP^ cKO mice at P40-60. Scale bars: 500 nm. **N-P**) Quantification of BSN (N; control: 100 ± 2.38 %; *n =* 8 mice; *atg5*^AgRP^ cKO: 92.73 ± 2.35 %; *n =* 8 mice), RIM-BP2 (O; control: 100 ± 6.33 %; *atg5^AgRP^* cKO:90.87 ± 1.93; *n =* 8 mice) spot density and RIM-BP2:BSN spot ratio (P; control: 0.9 ± 0.05; *atg5*^AgRP^ cKO: 0.9 ± 0.04; *n =* 8 mice/genotype) showed a significant decrease in BSN spot number/µm^2^ in *atg5*^AgRP^ cKO mice compared to controls (BSN spot density: p = 0.0475; *n =* 8 mice/genotype; Student’s t test). Circle size represents data belonging to one given animal (B, D, H). Graphs show median, lower and upper quartiles, whiskers represent min/max scores. *p<0.05, **p<0.01, ***p<0.001.

Our findings demonstrate that undisturbed autophagy in NPY/AgRP neurons is essential for maintaining appropriate NPY levels and likely its secretion, through non-cell autonomous mechanisms. Additionally, selective disruption of autophagy in hypothalamic NPY/AgRP neurons leads to SQSTM1/p62 accumulation in hippocampal hilar NPY-secreting neurons, suggesting a non-cell autonomous impairment of autophagy in these cells.

### The autophagy/NPY axis modulates presynaptic remodeling at hippocampal synapses

We previously showed that attenuating autophagy and NPY/sNPF signaling exclusively in the MB (via knockdown of ATG5) triggers presynaptic remodeling across the *Drosophila* brain, mimicking the aging phenotype[6,10]. We therefore asked whether neuronal autophagy in hypothalamic NPY/AgRP neurons regulates presynaptic remodeling via non-cell autonomous mechanisms also in mice. Our results indicate that the autophagic status of these neurons controls NPY and SQSTM1/p62 levels in hippocampal hilar NPY-secreting neurons in a non-cell autonomous manner. We therefore hypothesized that loss of hypothalamic neuronal autophagy, leading to non-cell autonomous alterations in hippocampal NPY signaling and autophagy, in turn might drive presynaptic remodeling. To this end, we used time-gated Stimulated Emission Depletion (gSTED) microscopy, a super-resolution technique achieving lateral resolution down to ∼40 nm, to analyze the density and size of presynaptic active zones (AZs). We labeled the AZ scaffold proteins Bassoon (BSN) and RIM-Binding Protein 2 (RIM-BP2) at hippocampal mossy fiber (MF-CA3) and Schaffer collateral (CA3-CA1) synapses in young *atg5*^AgRP^ cKO mice. These two synapse types exhibit marked differences in release probability and plasticity[21], with active zone remodeling possibly playing a major role in transmission at synapses executing plasticity presynaptically, e.g., MF-CA3 synapses. We found that *atg5*^AgRP^ cKO mice exhibited a significantly lower density of BSN spots at both synapse types (Figure 1I-P), while the ratio between RIM-BP2 and BSN spot density (RIM-BP2:BSN spot ratio; Figure 1L,P), the sheer expression levels of these two AZ components and their spot size did not significantly change (Fig. S2A-L).

We next asked whether NPY depletion alone is sufficient to drive presynaptic active zone (AZ) remodeling. To address this, we examined constitutive *npy* knockout (*npy* KO) mice[22] (Figure 2A-L) and analyzed their AZ status at the two synapse types introduced above. Notably, *npy* KO male mice showed a slight but statistically significant decrease in body weight between P40-P60 (Figure 2G,H), with no abnormalities. As we did not detect any sex-specific differences in NPY expression at this developmental stage (Figure 2A–F), male and female mice were pooled for subsequent analyses. At both CA3–CA1 and MF–CA3 synapses, NPY loss led to an increased density of RIM-BP2 spots or an elevated RIM-BP2:BSN spot ratio, respectively (Figure 2I–P), suggesting that NPY functions as a negative regulator of RIM-BP2 stabilization or recruitment at hippocampal presynaptic active zones. Our results suggest that changes in NPY signaling, similar to the observed shift after hypothalamic neuronal autophagy impairment, can regulate active zone remodeling at hippocampal synapses.

**Figure 2.**
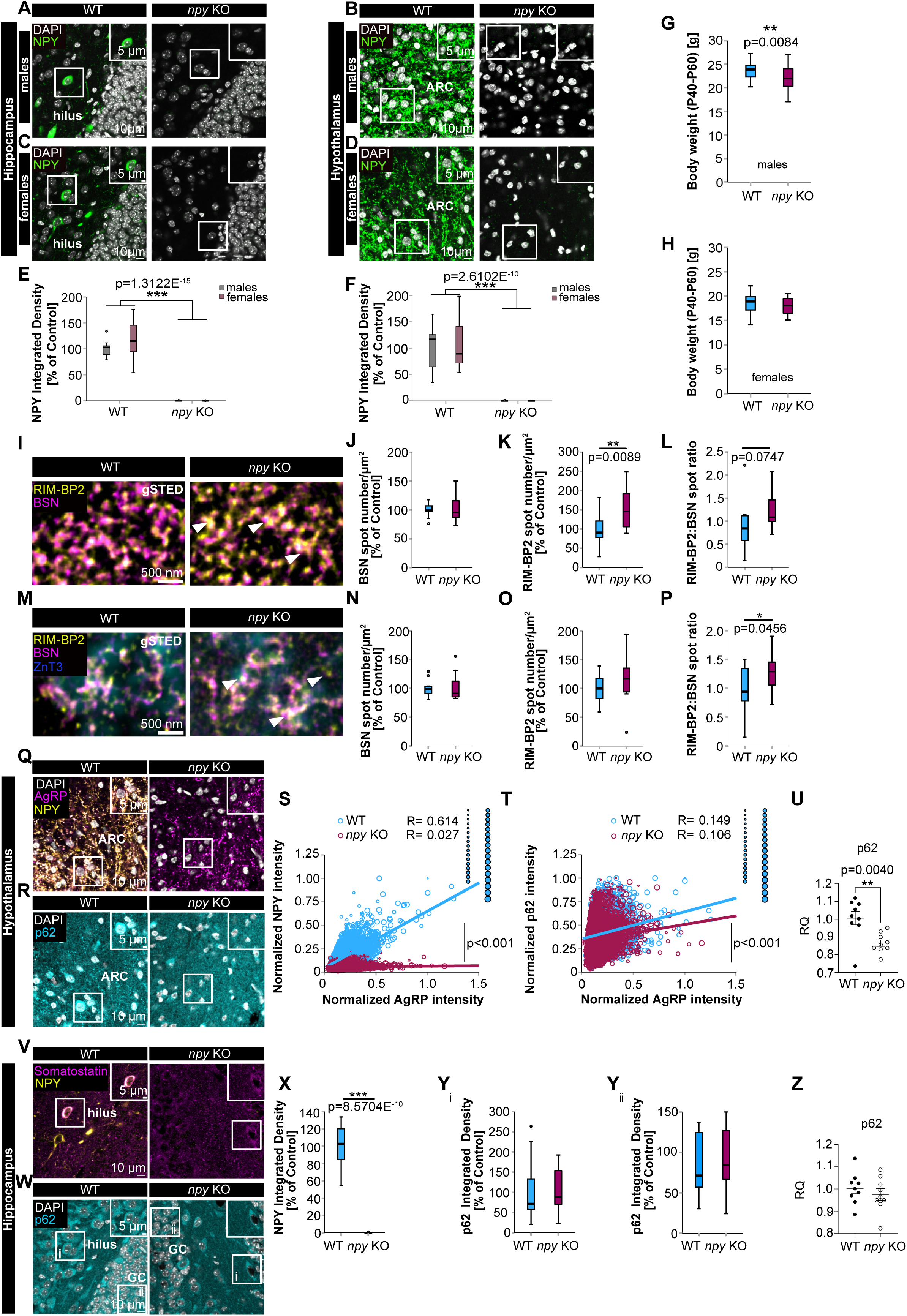
NPY negatively regulates hippocampal RIM-BP2 presynaptic recruitment while maintaining SQSTM1/p62 levels in the hypothalamus. **A-H**) Characterization of constitutive *npy* KO mice at P40-60. A) Representative confocal images of NPY immunolabelling in hippocampal and hypothalamic (B) NPY-secreting neurons of males WT and *npy* KO mice. C) Representative confocal images of NPY immunostaining in hippocampal and hypothalamic (D) NPY-secreting neurons of females WT and *npy* KO mice. Scale bars: 10 µm, insets: 5 µm. E) NPY integrated density of WT males (100 ± 5.57 %; *n* = 9 mice) and females (117.49 ± 13.9 %; *n* = 8 mice) compared to *npy* KO mice (males: 0.11 ± 0.09 %; *n =* 9 mice; females: 0.03 ± 0.01 %; *n =* 8 mice) within the hippocampal hilar region (genotype: F(1, 30) = 229.95; p = 1.3122E-15; sex: F(1, 30) = 1.47; p = 0.2341, two-way ANOVA). F) NPY integrated density of WT males (100 ± 14.25 %; *n =* 9 mice) and females (107.26 ± 17.36 %; *n =* 8 mice) compared to *npy* KO mice (males: 0.40 ± 0.17 %; *n =* 9 mice; females: 0.49 ± 0.22 %; *n =* 8 mice) within the hypothalamic ARC region (genotype: F(1, 30) = 85.88; p = 2.6102E-10; sex: F(1, 30) = 0.11; p = 0.7439; two-way ANOVA). G) Body weight comparison at P40-60 in males (WT: 23.63 ± 0.36 g; *n =*27 mice; *npy* KO: 22.12 ± 0.4 g; *n =* 36 mice) and females (H; WT: 18.66 ± 0.33 g; *n =* 32 mice; *npy* KO: 17.97 ± 0.33 g; *n =* 25 mice) WT and *npy* KO mice; constitutive NPY deletion significantly reduced body weight in males (p = 0.0084; Student’s t test). **I**) Time-gated STED microscopy of RIM-BP2 and BSN at hippocampal CA3-CA1 synapses in WT and *npy* KO mice at P40-60. Scale bars: 500 nm. **J-L**) Quantification of BSN spot density (J: WT: 100 ± 3.27 %; *n =* 12 mice; *npy* KO: 102.09 ± 6.08 %; *n =* 13 mice), RIM-BP2 spot density (K; WT: 100 ± 11.6 %; *n =* 12 mice; *npy* KO: 154.88 ± 15.01 %; *n =* 13 mice) and RIM-BP2:BSN spot ratio (L; WT: 0.89 ± 0.15; *n =* 12 mice; *npy* KO: 1.24 ± 0.12; *n =* 13 mice) showed increased RIM-BP2 spot density in *npy* KO compared to *WT* mice (RIM-BP2 spot density: p = 0.0089; Student’s t test. RIM-BP2:BSN spot ratio: p = 0.0747; Student’s t test). **M**) Time-gated STED microscopy of RIM-BP2 and BSN at hippocampal MF-CA3 synapses identified by their marked ZnT3 expression (confocal) in WT and *npy* KO mice at P40-60. Scale bars: 500 nm. **N**) Quantification of BSN spot density (WT: 100 ± 4.14 %; *n =* 12 mice; *npy* KO: 101.92 ± 6.32 %; *n =* 13 mice). **O**) Quantification of RIM-BP2 spot density (WT: 100 ± 6.64 %; *n =* 12 mice; *npy* KO: 116.7 ± 11.57 %; *n =* 13 mice) and RIM-BP2:BSN spot ratio (**P**; WT: 0.96 ± 0.12; *n =* 12 mice; *npy* KO: 1.28 ± 0.09; *n =* 13 mice) showed increased RIM-BP2:BSN spot ratio in *npy* KO compared to WT mice (RIM-BP2:BSN spot ratio: p = 0.0456; Student’s t test). **Q**) Representative confocal images of AgRP-NPY immunoreactivity in hypothalamic ARC of WT and *npy* KO mice at P40-60. Scale bars: 10 µm, insets: 5 µm. **R**) SQSTM1/p62 immunolabelling in the ARC of WT and *npy* KO mice. Scale bars: 10 µm, insets: 5 µm. **S**) NPY-AgRP mean intensity analysis in the hypothalamic ARC of WT and *npy* KO mice. NPY levels were related to AgRP levels with a GLMM with gamma regression (p < 0.001; WT: *n =* 6950 cells; npy KO: *n =* 7128 cells; *N* = 17 mice/genotype). Data obtained from male and female mice were pooled. **T**) SQSTM1/p62-AgRP mean intensity analysis showed that constitutive NPY deletion led to decreased SQSTM1/p62 expression in NPY/AgRP neurons. SQSTM1/p62 levels were related to AgRP levels with a GLMM with gamma regression (p < 0.001; WT: *n =* 6950 cells; npy KO: *n =* 7128 cells; *N =* 17 mice/genotype). **U**) RT-qPCR assay showed decreased SQSTM1/p62 transcript levels in the hypothalamus of *npy* KO mice at P40-60 (p = 0.0040; n = 9 mice/genotype; Mann-Whitney U-test; WT: 1.01 ± 0.04; *npy* KO: 0.87 ± 0.02). **V**) Representative confocal images of Somatostatin-NPY and SQSTM1/p62 (**W**) immunostaining in the dentate gyrus hilar region within the hippocampus of WT and *npy* KO mice at P40-60. Scale bars: 10 µm, insets: 5 µm. **X**) Hilar NPY Integrated density analysis (p = 8.5704E-10; WT: 100 ± 5.47 %; *npy* KO: 0.07 ± 0.05 %; *n =* 17 mice/genotype; Mann– Whitney U-test). **Yi and Yii**) Quantification of SQSTM1/p62 levels in the hippocampal hilar region (Yi; WT: 100 ± 16.57 %; *npy* KO: 107.43 ± 13.34 %; *n =* 17 mice/genotype) and granule cells layer (Yii; WT: 100 ± 15.25 %; *npy* KO: 100.11 ± 12.39 %; *n =* 17 mice/genotype) of WT and *npy* KO mice. Graphs show median, lower and upper quartiles, whiskers represent min/max scores. **Z**) SQSTM1/p62 transcript levels measured by RT-qPCR assay in the hippocampus of WT and *npy* KO mice at P40-60 (n = 9 mice/genotype; unpaired Student’s t test; WT: 1 ± 0.02; npy KO: 0.97 ± 0.03). Graphs show mean ± SEM (U and Z). Circle size represents data belonging to one given animal (S and T). *p<0.05, **p<0.01, ***p<0.001.

### NPY maintains SQSTM1/p62 transcription in hypothalamic NPY/AgRP neurons

Given the bidirectional cross-talk between neuronal autophagy and NPY, we assessed SQSTM1/p62 protein and transcript levels in both the hypothalamus and hippocampus of constitutive *npy* KO mice, using immunohistochemistry and RT-qPCR, respectively (Figure 2Q-Z). In *npy* KO mice, hypothalamic NPY/AgRP neurons exhibited significantly reduced SQSTM1/p62 levels compared to age-matched controls (Figure 2R,T). In contrast, SQSTM1/p62 protein levels in hippocampal NPY-expressing hilar neurons and granule cells were comparable in both groups, showing a high variability (Figure 2W,Yi,Yii). These observations were corroborated by RT-qPCR, which confirmed reduced *Sqstm1/p62* transcript levels in the hypothalamus (Figure 2U), suggesting that NPY supports *Sqstm1/p62* transcription in NPY/AgRP neurons, potentially influencing their autophagic activity or the recruitment of ubiquitinated cargos to autophagosomes.

### Dietary Spermidine supplementation increases NPY levels during aging

Several studies demonstrated Spd role in preventing senescence by activation of autophagy in aged cells, including neurons [6,9,23–25]. We previously showed that six months of dietary spermidine supplementation (Spd-S), initiated at 18 months of age, increased the levels of key autophagy-related proteins and reduced SQSTM1/p62 spot size in hippocampal neurons of aged mice, suggesting that chronic Spd administration may enhance autophagic activity in these neurons[26]. We therefore investigated whether spermidine supplementation (Spd-S), through activation of autophagy, could reverse age-related changes in NPY levels within the hippocampus and the hypothalamic arcuate nucleus (ARC), where NPY/AgRP neurons reside (Figure 3A-H).

**Figure 3.**
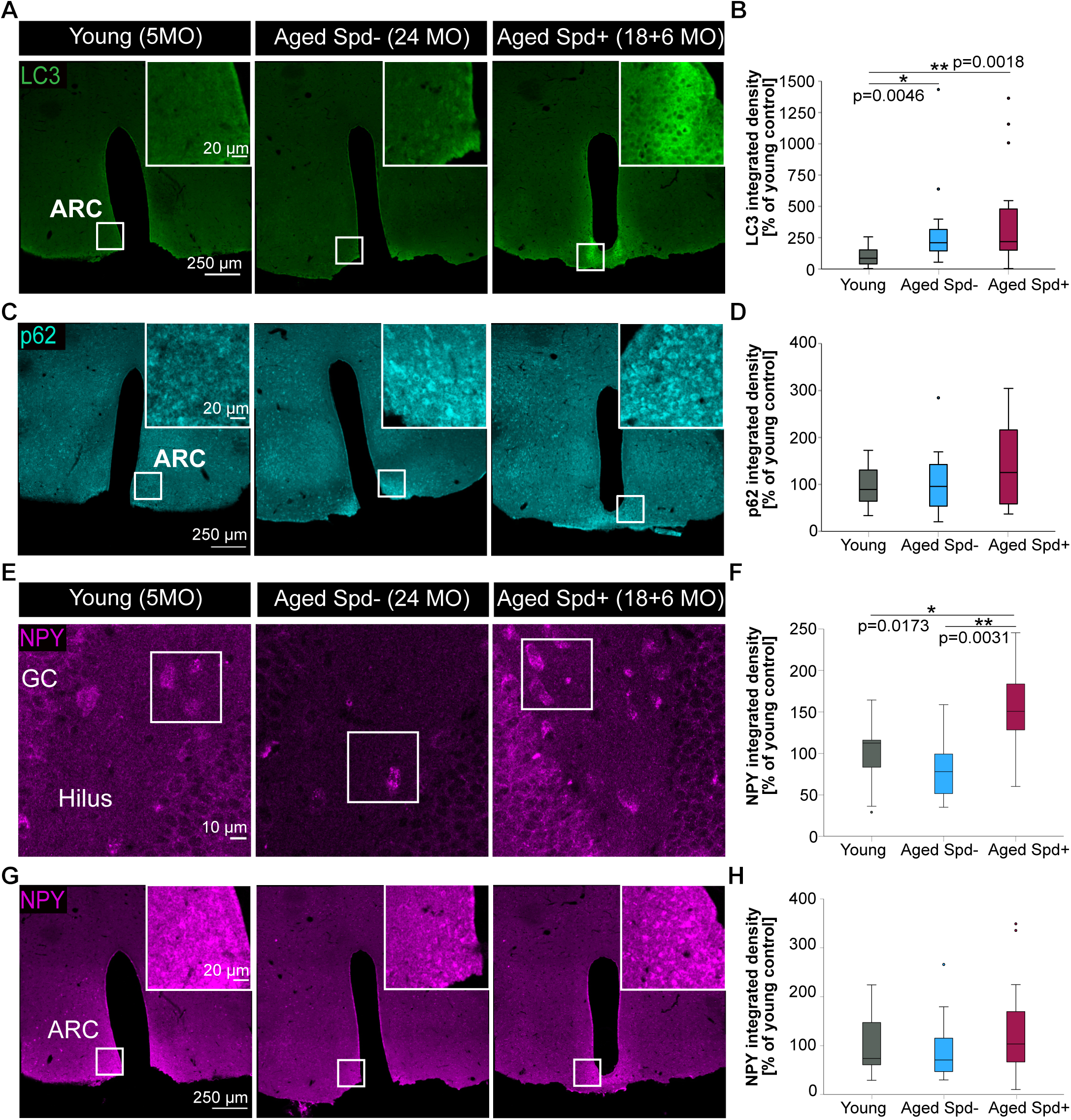
Spermidine supplementation protects NPY levels during aging. **A**) Representative confocal tile scans of LC3 immunoreactivity in hypothalamic ARC of young (5 months old), aged Spd-(24 months old) and aged Spd+ (18+6 months) treated mice. Scale bars: 250 µm, insets: 20 µm. **B**) Quantification of LC3 integrated density resulted in significantly increased LC3 levels in the hypothalamic ARC of aged Spd-(24MO; 301.97 ± 83.45 %; *n =* 16 mice) and aged Spd+ (18+6 MO; 395.47 ± 98.060 %; *n =* 17 mice) treated mice compared to young (5MO; 100 ± 18.83 %; *n =* 16 mice) controls (young versus aged Spd−: p = 0.0046; young versus aged Spd+: p = 0.0018; aged Spd− versus aged Spd+: p = 0.8062; p = 0.0026, Kruskal-Wallis test followed by Mann Whitney U test with Bonferroni correction setting α = 0.016667). **C**) Representative confocal tile scans of SQSTM1/p62 immunoreactivity in hypothalamic ARC of young (5MO), aged Spd-(24MO) and aged Spd+ (18+6 MO) treated mice. Scale bars: 250 µm, insets: 20 µm. **D**) Quantification of SQSTM1/p62 integrated density in the ARC of aged Spd-(24MO; 113.13 ± 34 %; *n =* 7 mice) and aged Spd+ (18+6 MO; 142.19 ± 26.3 %; *n =* 12 mice) treated animals compared to young (5MO; 100 ± 12.93 %; *n =* 13 mice) controls. **E**) NPY immunoreactivity in the hilar region of the dentate gyrus within the hippocampus of young, aged Spd-and aged Spd+ treated animals. Scale bar: 10 µm. **F**) Quantification of NPY levels within the hilar region of the hippocampus resulted in significantly increased NPY levels in aged Spd+ (18+6 MO; 152.8 ± 15.22 %; *n =* 12 mice) treated mice compared to young (5MO; 100 ± 11.26 %; *n =* 13 mice) and aged Spd-mice (24MO; 83.48 ± 13.02 %; *n =* 10 mice; F(2, 32) = 7.292; p = 0.0024, one-way ANOVA; young versus aged Spd+: p = 0.0173; aged Spd-versus aged Spd+: p = 0.0031). **G**) NPY immunoreactivity in the ARC with quantification of hypothalamic NPY levels (**H**) of young (5MO; 100 ± 16.52 %; *n =* 13 mice), aged Spd-(24MO; 91.21 ± 15.91 %; *n =* 16 mice) and aged Spd+ (18+6 MO; 133.62 ± 24.97 %; *n =* 16 mice) treated animals. Scale bars: 250 µm, insets: 20 µm. Graphs show median, lower and upper quartiles, whiskers represent min/max scores. *p<0.05, **p<0.01, ***p<0.001.

We first asked whether Spd-S enhances neuronal autophagy in the aged hypothalamic ARC, using immunohistochemistry. Aging was associated with a significant increase in overall LC3 levels, while SQSTM1/p62 expression remained unchanged (Figure 3A,B). Spd-S further elevated LC3 levels beyond those of young controls, again without significantly altering SQSTM1/p62 expression (Figure 3C,D). To evaluate the impact of aging and Spd-S on NPY expression, we assessed NPY levels in the hilus of the hippocampal dentate gyrus and the hypothalamic ARC across young, aged control, and aged Spd-S-treated mice. Aging led to a decline in hilar NPY expression, which was significantly restored, and even surpassed young levels, by Spd-S (Figure 3E,F). A similar, trend was observed in the ARC, but failing to reach significance (Figure 3G,H).

These findings suggest that Spd-S promotes autophagy and enhances NPY expression in aging NPY-expressing neurons of both the hippocampus and hypothalamus.

## Discussion

The decline of autophagy with age is increasingly recognized as a central contributor to neuronal dysfunction and synaptic deterioration[27–29]. Here, we identify a non-cell autonomous mechanism by which autophagy within hypothalamic NPY/AgRP neurons regulates presynaptic integrity in the hippocampus (Fig. S3). Conditional deletion of *atg5* in these neurons led to reduced NPY expression locally and distally, associated with presynaptic remodeling in hippocampal circuits, manifested as altered clustering of the key active zone protein BSN. These structural changes occurred without affecting overall AZ protein levels or spot size, suggesting selective disruption of AZ nanoarchitecture.

Constitutive *npy* knockout mice displayed similar remodeling phenotypes, implicating NPY as a downstream effector of hypothalamic autophagy. Additionally, NPY loss reduced SQSTM1/p62 expression in hypothalamic NPY/AgRP neurons, indicating a feedback loop in which NPY maintains autophagy-related gene expression. In contrast, hippocampal SQSTM1/p62 levels remained variable, hinting at region-specific regulation and possibly indirect effects.

Dietary Spd-S, known to induce autophagy and promote longevity[9,23,30], reversed aging-associated declines in NPY expression in both hypothalamus and hippocampus. This effect occurred alongside increased LC3 levels in the aged hypothalamus, suggesting restored autophagic activity. Notably, Spd-S elevated NPY expression even beyond young control levels, highlighting its potential to rejuvenate neuropeptide signaling.

Together, these findings suggest that hypothalamic autophagy, via NPY signaling, orchestrates presynaptic maintenance in distant brain regions. This pathway may integrate metabolic state with synaptic resilience and offers a promising target for dietary or pharmacological interventions aimed at preserving brain function during aging.

## Material and methods

### Animal experiments

All animal experiments were reviewed and approved by the Landesamt für Gesundheit und Soziales Berlin and by the animal welfare committee of Charité Universitätsmedizin Berlin, Leibniz Institut für Molekulare Pharmakologie (FMP) and Max-Delbrück-Center (MDC). All experiments were performed in accordance with the relevant guidelines and regulations. Mice were housed under 12/12-h light/dark cycle, with access to food and water ad libitum. *atg5^flox/flox^* [B6.129S-ATG5tm1Myok (RBRC02975)] mice[16] were received from the RIKEN BioResource Center (BRC, Ibaraki, Japan), AgRP-IRES-Cre [Agrptm1(cre)Lowl/J] mice[18] were purchased from the Jackson laboratories (Stock-Nr.: 012899). By mating AgRP-Cre;*atg5*^flox/+^ with *atg5*^flox/flox^ we obtained conditional AgRP-Cre;*atg5*^flox/flox^ (*atg5*^AgRP^ cKO) mice. To reduce the number of animals needed for experiments, both *atg5*^flox/flox^ and *atg5*^flox/+^ were used as control. *npy* KO mice in which the entire coding sequence including the initiation start codon was removed were previously reported[22]. Animals were investigated at postnatal day P40-60. Both males and females were used for all analyses. The number of animals required for this study has been determined by power analysis, based on own previous findings. Genotypes were determined by PCR analysis from genomic DNA prior experiments and after euthanasia. Primers to genotype mice belonging to the *atg5*^AgRP^ cKO strain were: Cre transgene (fw): 5′-GCGGTCTGGCAGTAAAAACTATC-3′; Cre transgene (rev): 5′-GTGAAACAGCATTGCTGTCACTT-3′; Cre internal positive control (fw): 5′-CTAGGCCACAGAATTGAAAGATCT-3′; Cre internal positive control (rev): 5′-GTAGGTGGAAATTCTAGCATCATC C-3′; to detect wild-type *Atg5* and *Atg5*^flox^ alleles[16,17]: A (fw): 5′-GAATATGAAGGCACACCCCTGAAATG-3′; B (rev): 5′-GTACTGCATAATGGTTTAACTCTTGC-3′; C (fw): 5′-ACAACGTCGAGCACAGCTGCGCAAGG-3′. *npy* KO mice were genotyped as previously described[22].

### Spermidine supplementation in mice

C57BL6 WT mice were purchased from Janvier Labs (C57BL/6 J: Rj males). Spermidine supplementation at a final concentration of 3 mM in drinking water started late in life (18 months of age) for 6 months, as previously described[26].

### Immunohistochemistry for confocal microscopy and image analysis

*npy* KO and *atg5*^AgRP^ cKO mice and respectively WT and control mice of both sexes (P40-60) were euthanized by an overdose (i.p.) of Pentobarbital (100-200 mg/kg body weight) and transcardially perfused with 1x PBS followed by 4 % (wt/vol) paraformaldehyde in 0.1 M PBS, pH 7.4. Mice belonging to Spd cohorts were euthanized by an overdose (i.p.) of Ketamin (120 mg/kg body weight)/Xylazin (16 mg/kg body weight) and transcardially perfused as above. Immunostainings on samples obtained from Spd cohorts were predominantly conducted on the same brain sections as previously published[26].

Immunostainings were performed as previously described[26], with antibodies listed in Table 1. Briefly, following permeabilization and blocking, 30 µm sections were first incubated in Fab fragment Donkey or Goat anti mouse IgG (H+L) (Jackson ImmunoResearch) for 1 h at room temperature (RT) and then with primary antibodies for 48 h at 4°C. Secondary antibody were applied for 2 h at RT, followed by washing and 20 min incubation with 2 μg/ml DAPI (Thermo Fisher Scientific, D1306). To quench lipofuscine autofluorescence, sections belonging to Spd cohorts were briefly rinsed with distilled water and incubated with 10 mM CuSO_4_ in 50 mM Ammonium Acetate (pH 5), 1 h at RT, rinsed with distilled water and finally washed 8 times for 5 minutes. Finally, slices were mounted with Mowiol pH 8 (Roth, 0713.1), using high precision coverslips No. 1.5H (Roth, LH25.2). Conventional confocal images were acquired with a Leica TCS SP8 confocal microscope (Leica Microsystems) equipped with a 63x, NA 1.4 oil immersion objective and a 20x, NA 0.75 oil immersion objective, for tile scan imaging. Sequential acquisition with Leica LAS X software (Leica Microsystems) was performed as previously described[26]. All settings were maintained equally for all groups within each experiment. Hypothalamic ARC and hippocampal regions were imaged within a given brain section. Experiments were performed two-four times on different biological replicates from three to four different mouse cohorts (Spd treatment) or from different litters. For the analysis, 8-bit images were thresholded and subsequently quantified by measuring the integrated density in a defined region of interest (ROI, 43.57 µm × 43.57μm or 81.2 µm x 81.2 µm for tile scans) using Fiji/ImageJ software (NIH). Within each round of experiments, the threshold was kept constant for all images of all groups within a given confocal channel. Data were then normalized to the mean of controls and pooled. For cell-based mean intensity measurements, a custom-made ImageJ macro was used (Kerkhoff Y. AgRP and ATG5 Quantification Macro. Zenodo; 2025. doi: 10.5281/zenodo.15464181). In short, DAPI labeled cell nuclei were Gaussian blurred with a sigma radius of 10, particles were then segmented upon thresholding with the find maxima function and the mean pixel intensity within the segmented region was quantified. Mean pixel intensities for a given channel were then normalized to the maximum of controls and pooled. Experiments and analysis were performed blindly.

**Table 1.** Antibodies for immunohistochemistry.

| Antigen | Species | Dilution | Source |
| --- | --- | --- | --- |
| AffiniPure Fab<br>Fragment Donkey Anti-Mouse IgG (H+L) | Donkey | 1:10 | Jackson<br>ImmunoResesarch; 715-007-003 |
| AffiniPure Fab<br>Fragment Goat Anti-Mouse IgG (H+L) | Goat | 1:10<br>1:25 (Spd cohorts) | Jackson<br>ImmunoResesarch; 115-007-003 |
| AgRP | Goat | 1:500 | Thermo Fisher Scientific;<br>PA5-47831 |
| ATG5 | Rabbit | 1:100 | LSBio; LS-C156610 |
| BSN | Mouse | 1:1200 | Abcam; ab82958 |
| LC3 | Mouse | 1:200 | MBL, M152-3 |
| NPY | Guinea Pig | 1:200 | Abcam; ab10341 |
| NPY | Rabbit | 1:500 | Cell Signaling; mAb 11976 |
| RIM-BP2 | Guinea Pig | 1:500 | Gentle gift by Dr. A.<br>Fejtova and Dr. E.<br>Gundelfinger |
| SQSTM1/p62 | Guinea Pig | 1:300 | Progen; GP62-C |
| SOMATOSTATIN-28 | Chicken | 1:300 | Synaptic Systems; 366 006 |
| ZnT3 | Rabbit | 1:500 | Synaptic Systems; 197 003 |
| Goat anti-guinea pig<br>Alexa Fluor 594 | Goat | 1:100 (gSTED)<br>1:400 (confocal) | Invitrogen; A-11076 |
| Goat anti-mouse STAR<br>RED | Goat | 1:300 | Abberior; STRED-1001-500UG |
| Goat anti-rabbit Alexa<br>Fluor 488 | Goat | 1:400 | Invitrogen; A-11034 |
| Donkey anti-chicken<br>Alexa Fuor 488 | Donkey | 1:200 | Jackson ImmunoResearch;<br>703-545-155 |
| Donkey anti-goat Alexa<br>Fluor 594 | Donkey | 1:200 | Jackson ImmunoResearch;<br>705-585-147 |
| Donkey anti-goat<br>ATTO490LS | Donkey | 1:100 | Hypermol; 2904-1MG |
| Donkey anti-guinea pig<br>Alexa Fluor 594 | Donkey | 1:400 | Jackson ImmunoResearch;<br>706-585-148 |
| Donkey anti-guinea pig<br>Alexa Fluor 647 | Donkey | 1:400 | Jackson ImmunoResearch;<br>706-605-148 |
| Donkey anti-guinea pig<br>Cy5 | Donkey | 1:200 | Jackson ImmunoResearch;<br>706-175-148 |
| Donkey anti-mouse<br>Alexa Fluor 488 | Donkey | 1:400 | Jackson ImmunoResearch;<br>715-545-151 |
| Donkey anti-rabbit<br>Alexa Fluor 488 | Donkey | 1:400 | Jackson ImmunoResearch;<br>711-545-152 |
| Donkey anti-rabbit<br>Alexa Fluor 594 | Donkey | 1:500 | Jackson ImmunoResearch;<br>711-585-152 |
| Donkey anti-rabbit<br>Alexa Fluor 647 | Donkey | 1:400 | Jackson ImmunoResearch;<br>711-605-152 |

**Table 2.**
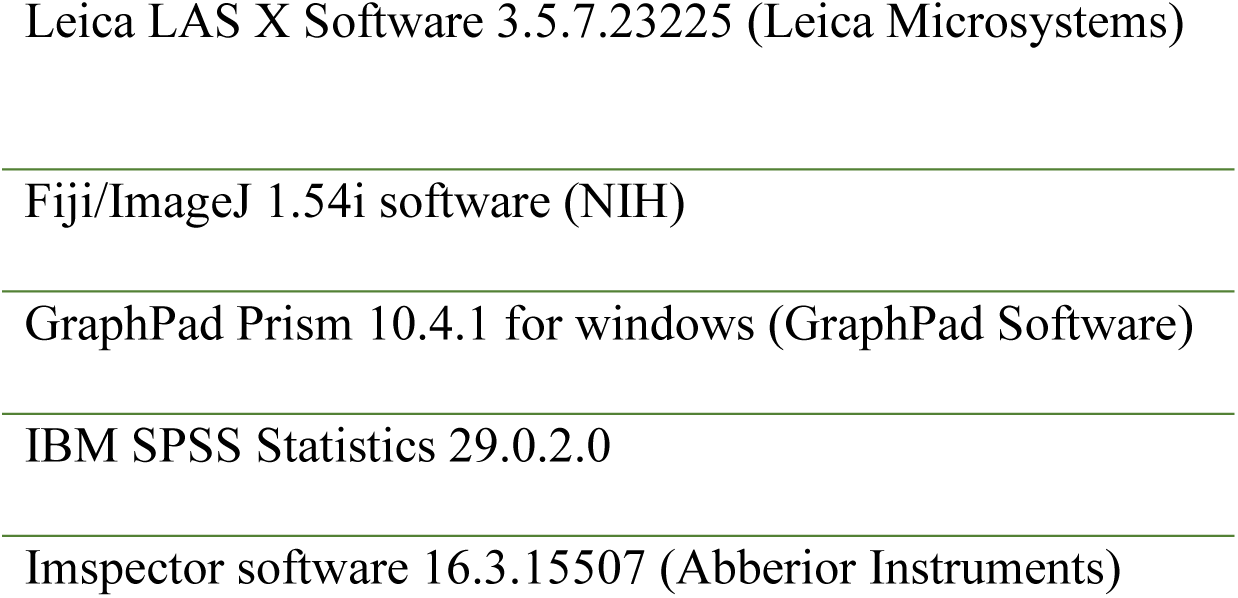
List of the software used for imaging and data analysis.

### Immunohistochemistry for time gated STED microscopy and image analysis

Brains were quickly removed from the skull and immediately shock-frozen on dry ice. The immunostaining was carried out as previously described[31], with antibodies listed in Table 1. Briefly, following fixation, 10 µm sagittal sections were blocked and permeabilized. Sections were first incubated with Fab fragment Goat anti mouse IgG (H+L) (Jackson ImmunoResearch) for 1 h at RT and then with primary antibody solution overnight in a humidified chamber at 4°C. Secondary antibodies were applied for 2 h at RT. Finally, sections were washed, dried and mounted on high precision coverslip No. 1.5H (Roth, LH25.2) with ProLong Gold (Cell Signaling Technology; 9071S).

Cryosections were imaged by time-gated STED (gSTED) microscopy using an Abberior Instruments STED Expert Line setup equipped with 100× oil-immersion objective lens (Olympus, NA = 1.4), as previously described[32]. Settings were maintained equal for sections of all groups within each experiment. Experiments and analysis were repeated at least two times on different biological replicates obtained from different litters. Raw gSTED images were deconvolved using a Richardson-Lucy algorithm (20 iterations) from Python’s scikit-image library with a bivariate Cauchy distribution as point-spread function with gamma set to 40 nm, as measured on crimson beads. Code has been made available via Github: https://github.com/KleistLab/STEDdeconvolution.

Segmented particle analysis was performed on deconvolved 8-bit and thresholded gSTED images with the analyze particles function in Fiji/ImageJ (NIH), as previously described[32]. For MF-CA3 synapses, BSN and RIM-BP2 spots were detected within the ZnT3 + area, creating a mask upon threshold on corresponding ZnT3 +confocal images[32]. Confocal images acquired in parallel to the raw gSTED images were used to measure the integrated density of a given channel upon signal thresholding. Within each round of experiments, the threshold was kept constant for all images of all groups within a given confocal channel. Data were then normalized to the mean of controls and pooled. Experiments and analysis were performed blindly.

### Reverse-transcriptase quantitative PCR assays (RT-qPCR)

RNA isolation from freshly dissected and snap-frozen brain tissue (hippocampus, hypothalamus) was done using the RNeasy Plus Mini kit (Qiagen, Cat. No. 74134) following manufacturer instructions. The RNA-eluate was then reverse transcribed into cDNA using the PrimeScript RT reagent kit (TaKaRa, RR037A). The reverse transcription was carried out in the Veriti 96-Well Thermocycler (Applied Biosystems) at 37°C for 15min, the subsequent inactivation was done at 85°C for 5min. The RT-qPCR was done using TaqMan Real-Time-PCR assays (ThermoFisher Scientific, p62 assay: Mm00448091_m1 Sqstm1) with the QuantStudio 3 (Applied Biosystems). GAPDH (Applied Biosystems, REF 4352339) was used as an endogenous control and as reference transcript.

RT-qPCR results were analyzed using the ΔCT method. First, the average was calculated for each group based on genotype and sex. This value was used to normalize the ΔCT of the respective group yielding the ΔΔCT which was then used for the relative quantification: RT = 2−(-ΔΔCT).

### Statistical analysis

Statistical analysis was performed with SPSS Statistics 29.0.2.0 (IBM). The statistical analysis of the RT-qPCR relative quantification was done using GraphPad Prism version 10.4.1 for windows (GraphPad Software, Boston, Massachusetts USA, www.graphpad.com). Data distribution was tested by using Shapiro-Wilk normality test and by checking histograms and normality Q-Q plots. Two groups dataset were compared using unpaired Student’s t test when normally distributed or non-parametric Mann–Whitney U-test for not normally distributed dataset. For comparisons with multiple groups, data were analyzed with one-way ANOVA followed by Tukey’s post-hoc test, or Kruskal–Wallis test followed by Mann Whitney U test with Bonferroni correction. Differences between groups for sex and genotype were analyzed with two-way ANOVA. Mendelian ratios were analyzed with one-sample Chi Square test. To account for nested data and fixed effects, a Generalized Linear Mixed Model (GLMM), with gamma regression was also used, having as fixed factor AgRP or NPY mean intensity. Linear correlation between mean intensities was determined with the Pearson correlation coefficient (R), reported in graphs. Only two tail p-values were considered. p-values, *n* and *N* are given in the figure legends. Values are represented as mean ± SEM. Asterisks denote significance: *p < 0.05; **p< 0.01; ***p < 0.001.

### Data availability

The data that supports the findings of this study are available from the corresponding author upon reasonable request.

## Abbreviations

AgRP: Agouti-related peptide
ATG5: autophagy related 5
AZ: active zone
BRP: Bruchpilot
Bsn: Bassoon
CA3-CA1: Cornu Ammmonis 3-Cornu Ammonis 1
cKO: conditional knockout
KO: knockout
LC3: microtubule associated protein 1 light chain 3
MB: Mushroom Body
MF-CA3: Mossy Fiber-Cornus Ammonis 3
NPY: neuropeptide Y
PreScale: presynaptic upscaling
RIM-BP2: RIM-Binding protein 2
SQSTM1/p62: sequestosome-1
sNPF: short Neuropeptide F
Spd: spermidine
Spd-S: spermidine supplementation

## Acknowledgements

The authors are grateful for the support by staff members of the animal facilities of the Leibniz Forschungsinstitut für Molekulare Pharmakologie (FMP), the Charité Universitätsmedizin and the Max-Delbrück-Center (MDC). We thank Silke Zillmann, Maria Mühlbauer, Anke Schönherr, Susanne Rieckmann and Anastasia Stawrakakis for technical assistance. We are grateful to Dr. Anna Fejtova and Dr. Eckart Gundelfinger for the RIM-BP2 antibody. We acknowledge the assistance of the core facility BioSupramol supported by the Deutsche Forschungsgemeinschaft (DFG, German Research Foundation) and the research center SupraFAB for the use and support of the Leica TCS SP8 confocal microscope and Abberior Instruments STED Expert Line. We thank the NeuroCure imaging facility for the use of the Leica TCS SP8 confocal microscope. The project is funded by the Deutsche Forschungsgemeinschaft (DFG, German Research Foundation) within the Research Unit 5228 Syntophagy (project ID 447288260) to AA (AL 2561/1-1), VH (HA2686/22-1), SJS (SI 849/15-1) and MM (MA 9763/1-1), the Research Unit 5289 to MvK and ER (project ID 453877723) and under Germany’s Excellence Strategy – Exc-2049-390688087 to DS, SJS, VH) and by the Bundesministerium für Bildung und Forschung to SJS (Smartage, 01GQ1420A), to DS (Smartage, 01GQ1420B) and to VH (Smartage, 01GQ1420C).

## Author contributions

GC, AA, SJS, MM designed research; GC, DT, G. Krause, G. Kochlamazashvili performed research; GC, DT, G. Krause, MM analyzed data; YK, ER software; JL, IMS, MM methodology; MvK, HH, DS, VH, SJS resources; ER, MvK, AA, DS, VH, SJS, MM acquired funding; SJS, MM writing - original draft.

