## Supplement for "Non-cell autonomous control of presynaptic remodeling by the hypothalamic autophagy/NPY axis"

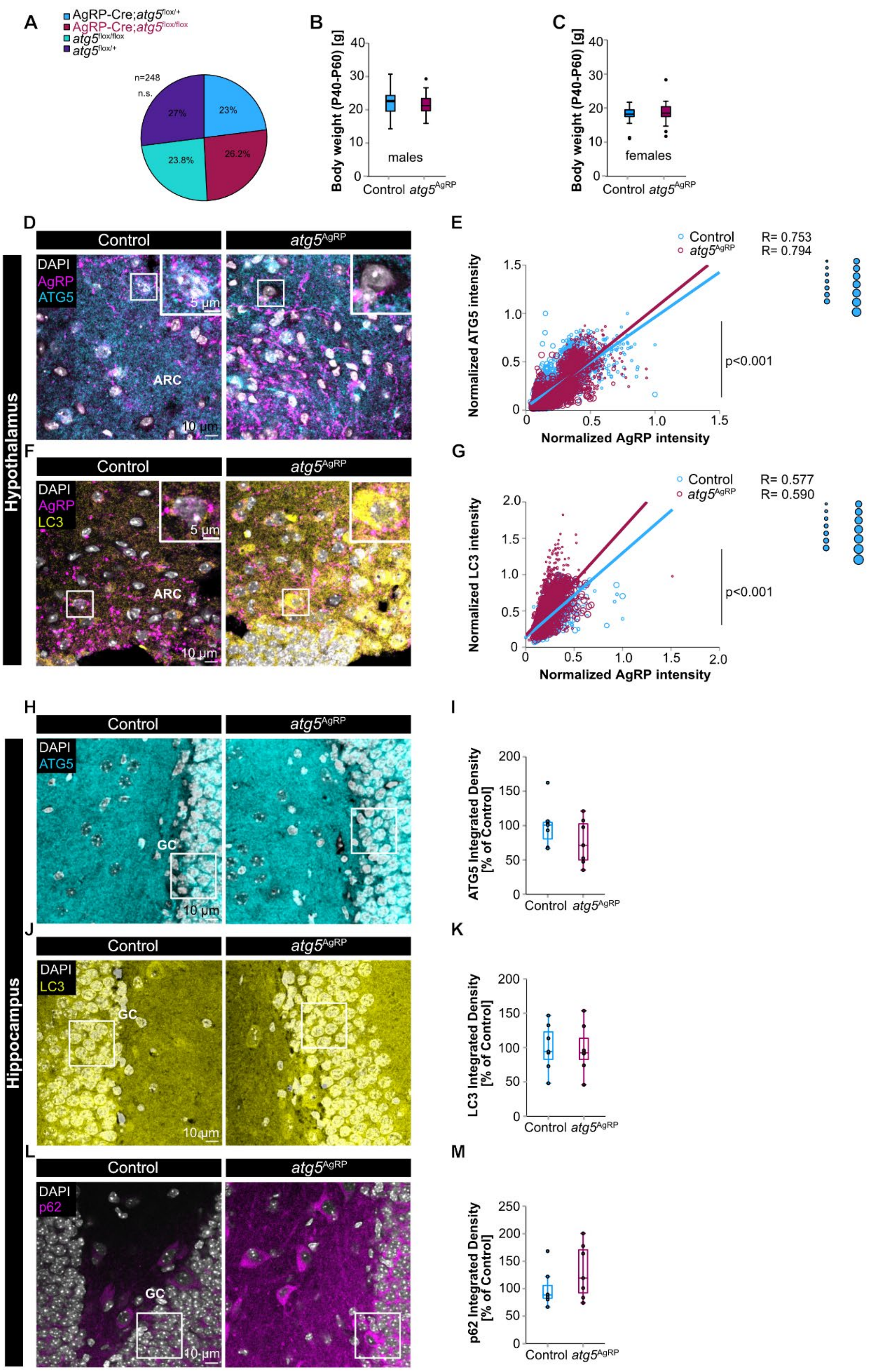


**Figure S1.** *atg5*$^{AgRP}$ cKO led to LC3 accumulation in the hypothalamic arcuate nucleus.

**A**) Mendelian distribution of *atg5*$^{\mathrm{AgRP}}$ progeny (*n* = 248 mice). **B and C**) Body weight comparison within P40-60 in males (B; control: 22.13 ± 0.55 g; *n* = 37 mice; *atg5*$^{\mathrm{AgRP}}$ cKO: 21.8 ± 0.57 g; *n* = 27 mice) and females (C; control: 18.15 ± 0.37 g; *n* = 42 mice; *atg5*$^{\mathrm{AgRP}}$ cKO: 18.66 ± 0.57 g; *n* = 29 mice) *atg5*$^{\mathrm{AgRP}}$ cKO and control mice. Graphs shows median, lower and upper quartiles, whiskers represent min/max scores. **D**) Representative confocal images of AgRP and ATG5 immunoreactivity in hypothalamic ARC of control and *atg5*$^{\mathrm{AgRP}}$ cKO mice at P40-60. **E**) ATG5-AgRP mean intensity analysis in the ARC of control and *atg5*$^{\mathrm{AgRP}}$ cKO mice. ATG5 levels were related to AgRP levels with a GLMM with gamma regression ($p < 0.001$; control: *n* = 1873 cells; *atg5*$^{\mathrm{AgRP}}$ cKO: *n* = 2150 cells; *N* = 7 mice/genotype). **F**) AgRP and LC3 immunolabelling in hypothalamic ARC of control and *atg5*$^{\mathrm{AgRP}}$ cKO mice at P40-60. **G**) LC3-AgRP mean intensity analysis showed an increase in the autophagosomal marker LC3 in hypothalamic NPY/AgRP neurons of *atg5*$^{\mathrm{AgRP}}$ cKO mice compared to controls. LC3 levels were related to AgRP levels with a GLMM with gamma regression ($p < 0.001$; control: *n* = 2791 cells; *atg5*$^{\mathrm{AgRP}}$ cKO: *n* = 2728 cells; *N* = 7 mice/genotype). Circle size represents data belonging to one given animal (E and G). **H**) Hippocampal granule cell layer immunostained for ATG5, followed by quantification of ATG5 levels (**I**) in control (100 ± 12.06 %; *n* = 7 mice) and *atg5*$^{\mathrm{AgRP}}$ cKO (76 ± 12.48 %; *n* = 7 mice). **J**) Hippocampal granule cell layer immunostained for LC3, followed by quantification of LC3 levels (**K**) in control (100 ± 12.85 %; *n* = 7 mice) and *atg5*$^{\mathrm{AgRP}}$ cKO (97.75 ± 13.42 %; *n* = 7 mice). **L**) Hippocampal granule cell layer immunostained for p62, followed by quantification p62 levels (**M**) in control (100 ± 10.09 %, *n* = 9 mice) and *atg5*$^{\mathrm{AgRP}}$ cKO mice (119.36 ±.16.71 %, *n* = 9 mice). Scale bars: 10 µm, insets: 5 µm. Graphs show median, lower and upper quartiles, whiskers represent min/max scores. *p<0.05; **p<0.01 ***p<0.01.

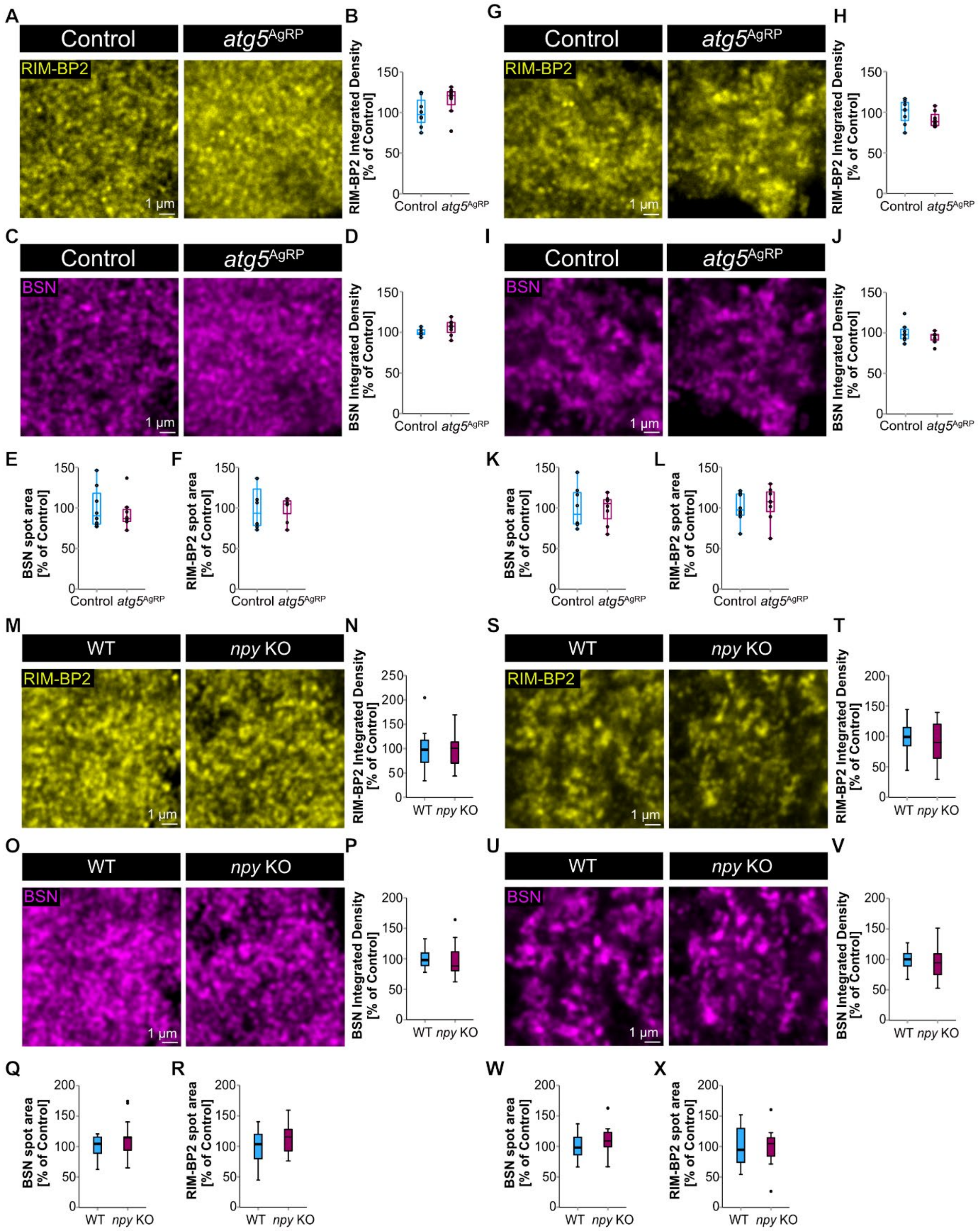


**Figure S2.** The autophagy/NPY axis does not significantly alter either BSN and RIM-BP2

levels or spot size at hippocampal synapses.

**A**) Representative confocal images of hippocampal CA3-CA1 synapses immunostained for RIM-BP2 in control and *atg5*$^{AgRP}$ cKO mice at P40-60, followed by RIM-BP2 (**B**) integrated density quantification (control: 100 ± 6.29 %; *atg5*$^{AgRP}$ cKO: 115 ± 6.24 %; *n* = 8 mice). **C**) Representative confocal images of hippocampal CA3-CA1 synapses immunostained for BSN in control and *atg5*$^{AgRP}$ c*KO* mice, followed by (**D**) BSN integrated density quantification (control: 100 ± 1.49 %; *atg5*$^{AgRP}$ cKO: 106 ± 3.31 %; *n* = 8 mice). **E**) Quantification of BSN and RIM-BP2 (**F**) spot area acquired by gSTED (see Figure 1) in control (BSN spot area: 100 ± 8.97 %; RIM-BP2 spot area: 100 ± 9.28 %; *n* = 8 mice) and *atg5*$^{AgRP}$ cKO mice (BSN spot area %: 93.17 ± 6.9; RIM-BP2 spot area:.105.57 ± 8.96 %; *n* = 8 mice). **G**) Representative confocal images of hippocampal MF-CA3 synapses immunostained for RIM-BP2 in control and *atg5*$^{AgRP}$ cKO mice at P40-60, followed by RIM-BP2 (**H**) integrated density quantification (control: 100 ± 5.16 %; *atg5*$^{AgRP}$ cKO: 91.07 ±3.32 %; *n* = 8 mice/genotype). **I**) Representative confocal images of hippocampal MF-CA3 synapses immunostained for BSN in control and *atg5*$^{AgRP}$ cKO mice at P40-60, followed by (**J**) BSN integrated density quantification (control: 100 ± 4.05 %; *atg5*$^{AgRP}$ cKO: 94.54 ± 2.45 %; *n* = 8 mice/genotype). Quantification of BSN (**K**) and RIM-BP2 (**L**) spot area acquired by gSTED (see Figure 1) in control (BSN spot area: 100 ± 8.91 %; RIM-BP2 spot area: 100 ± 6.33 %; *n* = 8 mice) and *atg5*$^{AgRP}$ cKO mice (BSN spot area: 98.76 ± 6.33 %; RIM-BP2 spot area: 104.52 ± 7.47 %; *n* = 8 mice). **M**) Representative confocal images of hippocampal CA3-CA1 synapses immunostained for RIM-BP2 in WT and *npy* KO mice at P40-60, followed by (**N**) RIM-BP2 integrated density quantification (WT: 100 ± 12.75 %; *n* = 12 mice; *npy* KO: 99.76 ± 11.04 %; *n* = 13 mice). **O**) Representative confocal images of hippocampal CA3-CA1 synapses immunostained for BSN in WT and *npy* KO mice at P40-60, followed by (**P**) BSN integrated density quantification (WT: 100 ± 4.78 %; *n* = 12 mice; *npy* KO: 97.46 ± 7.88 %; *n* = 13 mice). **Q**) Quantification of BSN spot area (WT: 100 ± 5.27 %; *n* = 12 mice; *npy* KO: 115.06 ± 8.68 %; *n* = 13 mice) and RIM-BP2 spot area (**R**; WT: 100 ± 8.19 %; *n* = 12 mice; *npy* KO: 114.87 ± 7.35 %; *n* = 13 mice) on gSTED images (see Figure 2) in WT and *npy* KO mice at P40-60. **S**) Representative confocal images of hippocampal mossy fiber-CA3 synapses immunostained for RIM-BP2 in WT and *npy* KO mice at P40-60, followed by (**T**) RIM-BP2 integrated density quantification (WT: 100 ± 8.22 %; *n* = 12 mice; *npy* KO: 91.3 ± 10.04 %; *n* = 13 mice). **U**) Representative confocal images of hippocampal MF-CA3 synapses immunostained for BSN in WT and *npy* KO mice at P40-60, followed by (**V**) BSN integrated density quantification (WT: 100 ± 5 %; *n* = 12 mice; *npy* KO: 95.91 ± 7.61 %; *n* = 13 mice). **W**) Quantification of BSN spot area (WT: 100 ± 5.73 %; *n* = 12 mice; *npy* KO: 111.39 ± 8.17 %; *n* = 13 mice) and RIM-BP2 spot area (**X;** WT: 100 ± 9.4 %; *n* = 12 mice; *npy* KO: 99.06 ± 8.84 %; *n* = 13 mice) on gSTED images (see Figure 2) in WT and *npy* KO mice. Scale bars: 1 µm. Graphs show median, lower and upper quartiles, whiskers represent min/max scores.

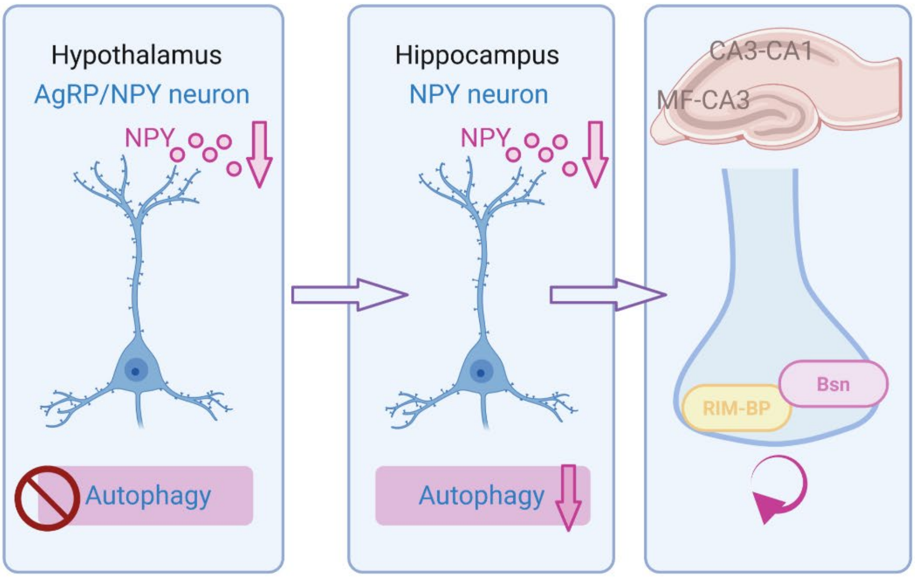


**Figure S3.** Non-cell autonomous regulation of hippocampal autophagy and presynaptic remodeling upon *atg5*$^{AgRP}$ cKO. Created in BioRender. Maglione, M. (2025) https://BioRender.com/9kcn9qj
